# Integrating Virtual Pivot Point and Trunk Dynamics to Understand Human Walking on Slopes: Insights from Experiments and Modeling

**DOI:** 10.64898/2026.02.23.707466

**Authors:** Vahid Firouzi, Johanna Vielemeyer, Andre Seyfarth, Oskar von Stryk, Roy Müller

**Affiliations:** Computer Science Department, TU Darmstadt, Darmstadt, Germany; GaitLab, Klinikum Bayreuth GmbH, Bayreuth, Germany; Friedrich Schiller University Jena, Jena, Germany; Centre for Cognitive Science, TU Darmstadt, Darmstadt, Germany; BaySpo, University of Bayreuth, Germany; Friedrich-Alexander-University Erlangen-Nürnberg, Germany

## Abstract

Walking on sloped terrain requires substantial mechanical and control adaptations for effective energy management compared to level ground locomotion. The Virtual Pivot Point (VPP) hypothesis explains sagittal plane angular momentum regulation during level walking, but its validity in slope walking remains unexplored. This study combines human experiments with template-model simulations to investigate how the VPP strategy is modulated during slope walking. Participants walked on an instrumented ramp at various inclinations (0°, ± 7.5°, ± 10°), while a 2D spring-loaded inverted-pendulum model with a trunk segment simulated the task. Experimental results confirmed that the VPP is a robust feature of slope walking (*R*^2^ > 0.975). The simulation reproduced the change in hip torque and trunk adaptations by modulating VPP position. Results of this study indicate that VPP position and trunk dynamics could afford stability and energy management on gentle slopes, but to robustly navigate steeper ramps, humans recruit a multi-joint strategy where the knee and ankle joints play a crucial role in managing the energetic demands of sloped terrain. Beyond advancing our understanding of locomotor control, these insights have practical implications for the design of exoskeletons that adapt to uneven terrain.

## 1 Introduction

Walking on varied terrain is a hallmark of human bipedalism, yet it presents significant challenges to the neuromuscular system. Navigating slopes, in particular, requires sophisticated adjustments to maintain stability and manage the body’s mechanical energy. Ascending a slope demands positive work to propel the body forward and upward, while descending requires negative work to control the descent and safely dissipate energy [1, 2]. Understanding the underlying control strategies that enable these adaptations is crucial not only for fundamental biomechanics but also for developing effective assistive technologies and rehabilitation protocols.

A compelling model for explaining the control of upright walking is the Virtual Pivot Point (VPP) hypothesis [3, 4]. This concept posits that the ground reaction forces (GRFs) converge at a point located above the body’s center of mass (CoM) in the sagittal plane during the single support phase of walking. In healthy adults, the consistent use of a VPP strategy is fundamental to regulating whole-body angular momentum in the sagittal plane during level-ground walking, a finding that holds true across numerous conditions, such as walking barefoot, with shoes, with added mass, or with flexed hip [3, 5–7]. The VPP framework has also been successfully applied to analyze more dynamic gaits, such as running [8]. Recent work has highlighted the VPP’s utility in explaining age-related gait adaptations and instability in Parkinsonian gait [4, 9] on level ground walking. This established robustness on level ground suggests the VPP is a core feature of human gait control, yet it simultaneously raises a critical question: is this strategy preserved when the mechanical demands of walking are fundamentally altered?

Walking on sloped terrain provides a natural and challenging test bed for this very question, as the constant change in the gravitational vector fundamentally shifts the mechanical and energetic demands compared to level walking [10–12]. These altered requirements—from the net positive mechanical work needed uphill to the controlled energy dissipation required downhill [1, 13]—directly necessitate significant adaptations in gait kinematics, kinetics, and, crucially, the regulation of angular momentum [11, 14]. For instance, downhill walking is associated with tighter momentum control to mitigate fall risk, while uphill walking allows for greater momentum excursions to meet higher propulsive demands [11, 14]. These findings, therefore, imply that the VPP strategy, if applicable to slope walking, must also adapt to the asymmetrical challenges of slope walking.

This study combines experimental analysis with template-model simulations to investigate VPP dynamics and the associated control mechanisms during uphill and downhill walking. Our objectives are to: (i) establish the validity of the VPP concept across varying slopes and quantify its location and consistency; (ii) analyze the crucial kinematic and kinetic adaptations required at the lower limb joints to achieve the VPP strategy on slopes, particularly focusing on their modulation of torque and power; and (iii) determine the integrated role of trunk dynamics in achieving the VPP control strategy, proposing a VPP-informed template model that successfully reproduces the observed slope-specific adaptations. By unifying experimental data with a predictive modeling framework, this work clarifies the VPP’s function in modulating energy on slope, offering foundational insights for robotics, prosthetics, and rehabilitative science.

## 2 Methods

In this section, we first describe the experimental protocol and data acquisition procedures used to capture the biomechanical measurements. We then detail the data processing and analysis methods applied to the experimental data. Finally, we present the simulation framework and control strategy used to model the walking on the slope.

### 2.1 Experimental Methods

#### 2.1.1 Participants

In this study, 13 able-bodied young adults (five women, eight men) with a mean age of 28.2 ± 4.7 years, mean body mass of 73.3 ± 12.4 kg and mean height of 1.78 ± 0.09 m took part. Three of the participants were excluded in the analysis because of missing data. The protocol was approved by the ethics committee of the University of Jena (5368-12/17) and was conducted in accordance with the Declaration of Helsinki. Prior to experiments, the participants signed an informed consent form.

#### 2.1.2 Measurements

Participants walked up and down a 6 m instrumented ramp at inclinations of 0°, 7.5°, and 10° at self-selected speed (see Fig. 1b). Three force plates (AMTI Optima BMS464508) were positioned in the middle of the ramp and recorded ground reaction forces. Kinetic data were sampled at 1000 Hz. Kinematic data were measured with ten infrared cameras (Vero v2.2) and two video cameras (Vue) of Vicon (200 Hz). The subjects were prepared for 3D gait analysis following the standard protocol for the Plug-in Gait full body marker set [15].

**Figure 1.**
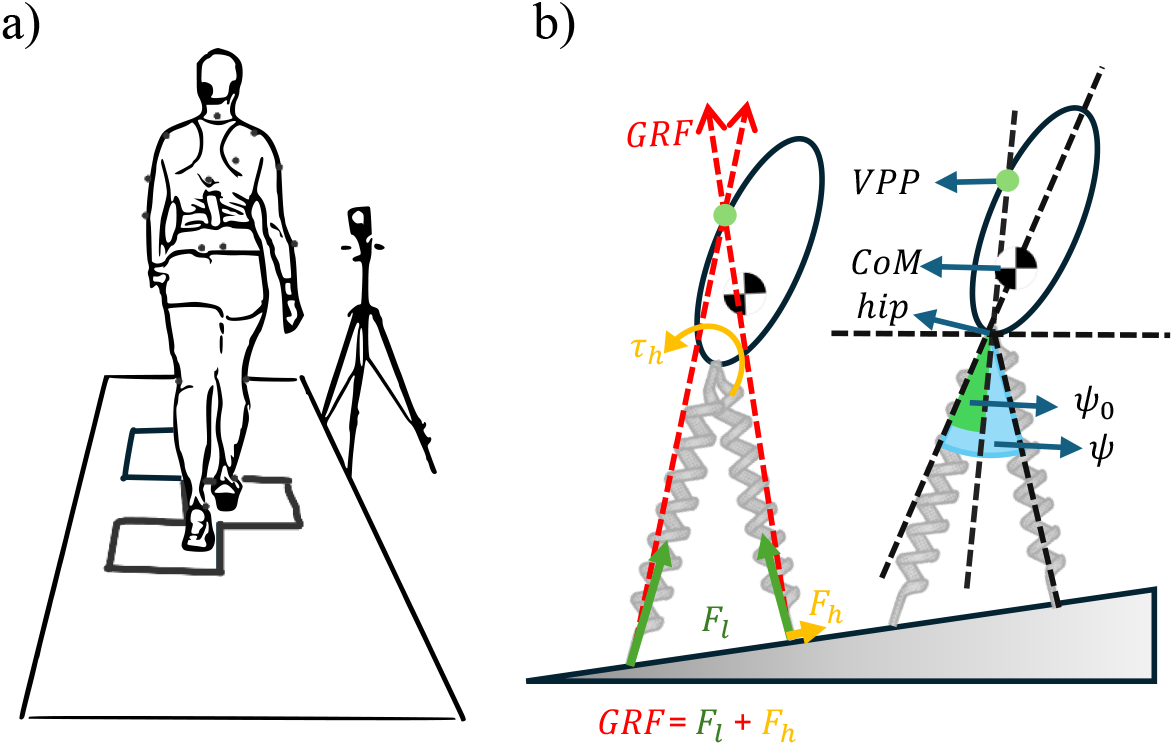
a) Experimental setup. b) The spring-loaded inverted pendulum (SLIP) model with a trunk walking on the slope.

For each condition, twelve valid trials were recorded after a few times walking up and down the ramp to familiarize themselves with the inclination. Participants were not informed about the force plates’ location to avoid targeting. A trial was considered valid when each foot made a clean contact on a separate force plate without overstepping. Ramp inclinations were tested in a randomized order across participants.

#### 2.1.3 Data processing

The raw data were processed in Nexus 2.16.0×64 (Vicon Motion Systems Ltd). For labeled trajectories, gap filling and filtering (mean square error setting) based on Woltring [16] was used. For calculation of all biomechanical parameters the Plug-in Gait full body model was run. Then, model data were exported as c3d and txt. This data were analyzed with custom-written Python code.

For GRFs, only two of three consecutive footfalls were analyzed to increase the rate of clean contacts. To compute the VPP, GRF vectors were used, starting in the center of pressure (CoP) for every instant of measurement. The position of the VPP was calculated as the point where the sum of the squared perpendicular distances to the GRFs is minimal. It was considered in a coordinate frame centered on the CoM and aligned to the trunk. The amount of agreement between model forces (vectors directed from the CoP to the VPP) and GRFs are represented by the coefficient of determination *R*^2^ [5, 17]. To compare VPP variables between level walking and ramp walking, t-tests (*P* < 0.05) were used. To identify differences in trunk angle, joint moments and joint power between level and ramp walking, we performed a point-by-point t-test across the normalized gait cycle. This analysis produced a set of p-values across the gait cycle (100 time points × 4 slopes). To control the Type I error rate, false discovery rate (FDR) correction was applied [18]. Significant intervals were defined using a predefined criterion: any segment with at least three consecutive time points showing p-values below 0.05 was considered a significant deviation from level walking. These significant intervals are illustrated in Figure 3 as bars above/below the data curves. Note that no significant differences were found for the trunk angle. Mean values over all trials were calculated to describe differences between conditions. A more detailed description of the measurements and data processing of the underlying data set can be found in a data descriptor paper [19].

### 2.2 Simulation Methods

To investigate the role of trunk dynamics in walking on the slope, we extended the traditional 2D spring-loaded inverted pendulum (SLIP) model [20] by incorporating a trunk segment attached to the hip joint. To regulate the trunk’s motion, we employed a Force Modulated Compliant Hip (FMCH) controller [4, 21], a bioinspired approach that dynamically adjusts hip joint stiffness based on leg force feedback. This controller modulates the hip torque to maintain trunk stability. The FMCH controller is designed to emulate the VPP hypothesis, which describes how GRFs converge at a virtual point above the CoM in a reference frame attached to the CoM and aligned with upper body orientation [3, 22]. The hip torque generated by the FMCH controller follows this equation:

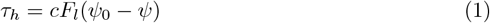

where *τ*_*h*_ represents the hip torque, *c* is a constant, *F*_*l*_ denotes the leg force, *ψ* is the angle between the upper body and the leg, and *ψ*_0_ is the rest angle, defined as the angle between the upper body orientation and the hip-VPP line (see Fig. 1b).

To investigate the role of trunk dynamics in slope walking, we performed an optimization process using Bayesian optimization to determine the model parameters and initial conditions necessary for stable locomotion on a ±1° incline [23].

## 3 Results

### 3.1 Experimental Results

The experimental results are illustrated in Figures 2, 3 and 4 and listed in Table 1. Additionally, significant mean differences will be reported in the following.

**Table 1.** Virtual pivot point (VPP; trunk aligned, center of mass (CoM) centered) and trunk values. Mean values ± standard deviation over all trials, participants and two consecutive contacts (single support phase, VPP values) or gait cycle (trunk values). Horizontal position is denoted by ‘x’, vertical by ‘z’, ‘osc.’ means ‘oscillation’. VPPz, *R*^2^ and mean trunk angle show significant differences between conditions. Aterisk (*) denotes significant differences from level walking.

|  | ramp 10° up | ramp 7.5° up | level | ramp 7.5° down | ramp 10° down |
| --- | --- | --- | --- | --- | --- |
| VPPx [m] | -0.040±0.026 | -0.048±0.034 | -0.039±0.013 | -0.032±0.017 | -0.036±0.020 |
| VPPz [m] | 0.503±0.108* | 0.477±0.127* | 0.307±0.086 | 0.206±0.075* | 0.219±0.092 |
| $R^2$ | 0.981±0.009* | 0.975±0.020* | 0.994±0.003 | 0.989±0.008 | 0.984±0.0011* |
| trunk [°] | 6.9±4.9* | 4.9±5.6 | -0.2±4.8 | -1.5±5.3 | -1.9±4.9 |
| trunk osc. [°]<br>(min/max) | 2.4 (6.1/8.5) | 2.6 (3.9/6.5) | 2.6 (-1.6/1.0) | 1.9 (-2.5/-0.6) | 1.6* (-2.8/-1.1) |

**Figure 2.**
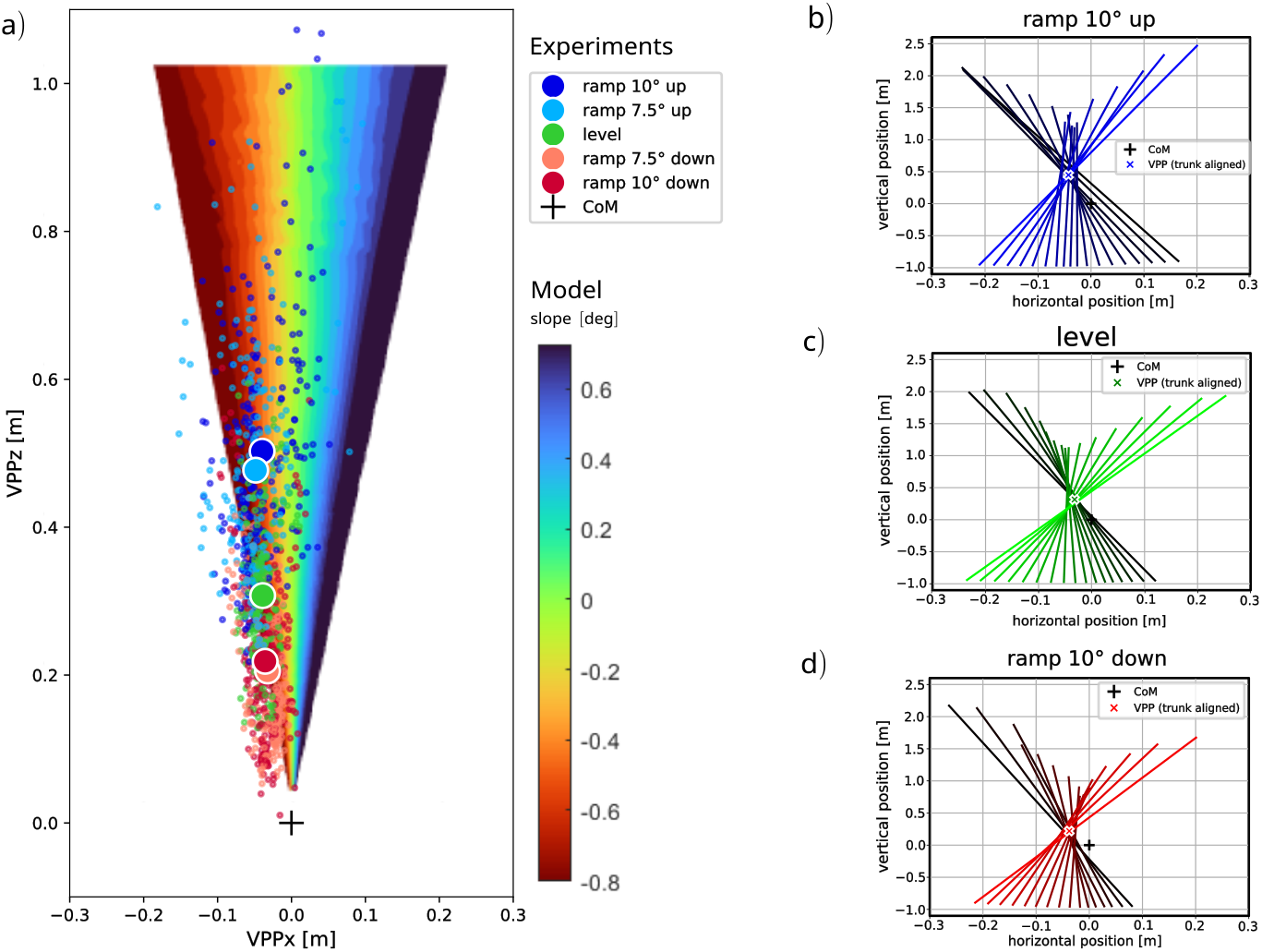
a) Experimental and model results of virtual pivot point (VPP) calculation. Small dots represent all trials of all participants, big circles illustrate the mean values over trials and participants, separate for each ramp condition. b)-d) Exemplarily VPP plots for single representative trials of one participant. Colored lines show the ground reaction forces (GRFs) at different measurement times originating at the center of pressure in a coordinate system centered on the center of mass (CoM) and aligned with the trunk. The GRFs are illustrated from touchdown (black line) to take-off (colored line). Colored crosses indicate the calculated VPP, black plus illustrates the CoM (zero point). b) walking ramp of 10° upwards, c) level walking, d) walking ramp of 10° downwards.

**Figure 3.**
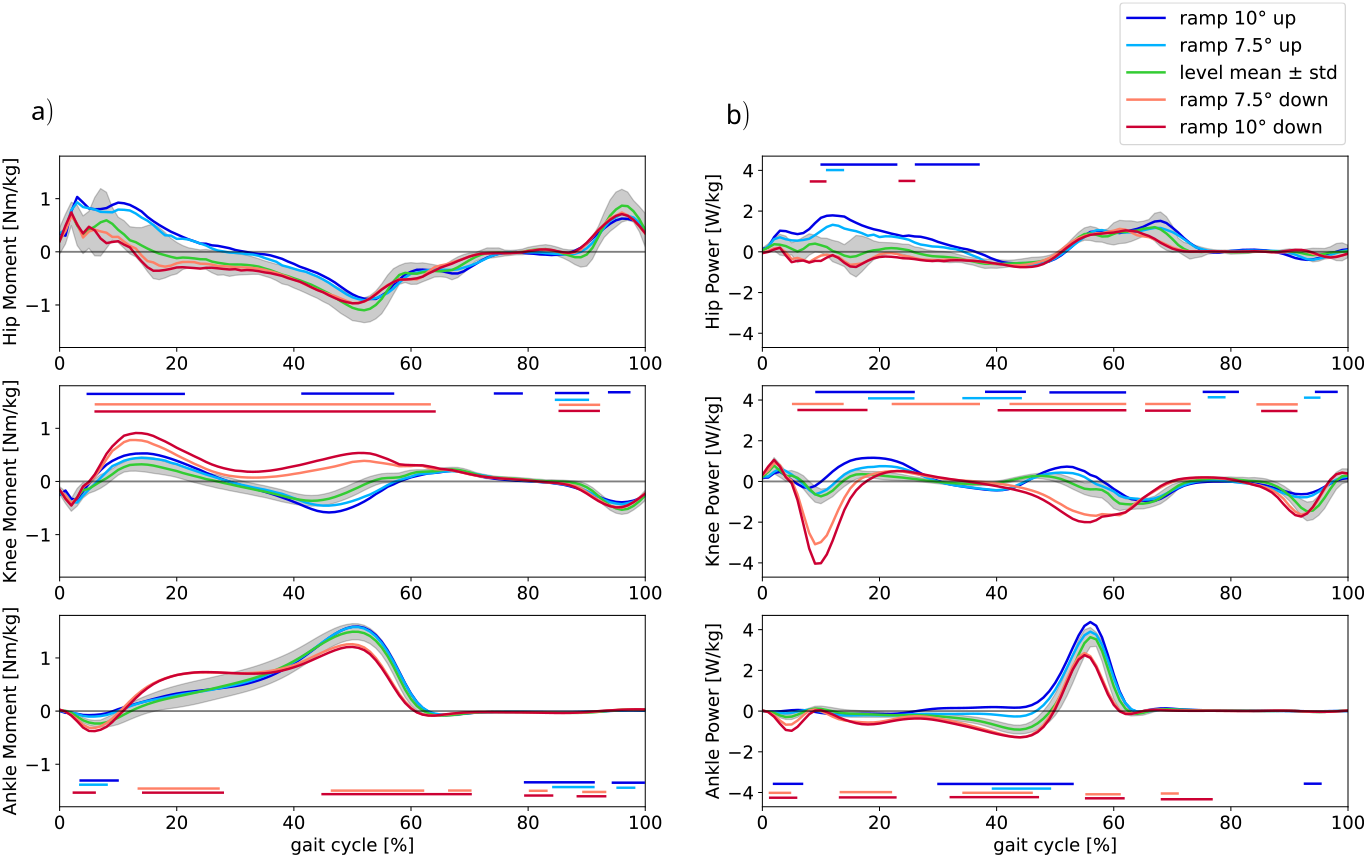
Joint moments (a) and power (b) of hip, knee and ankle joint of human experiments. Mean values over all trials and participants are shown. The gait cycle starts with touch down of the ipsilateral leg and ends with the next touch down of this leg. For knee and ankle, this ipsilateral leg is analyzed. For level walking, also standard deviation is shown (gray shaded area). Bars show significant differences from level walking, it is illustrated if it was significant for at least three consecutive measurement points.

**Figure 4.**
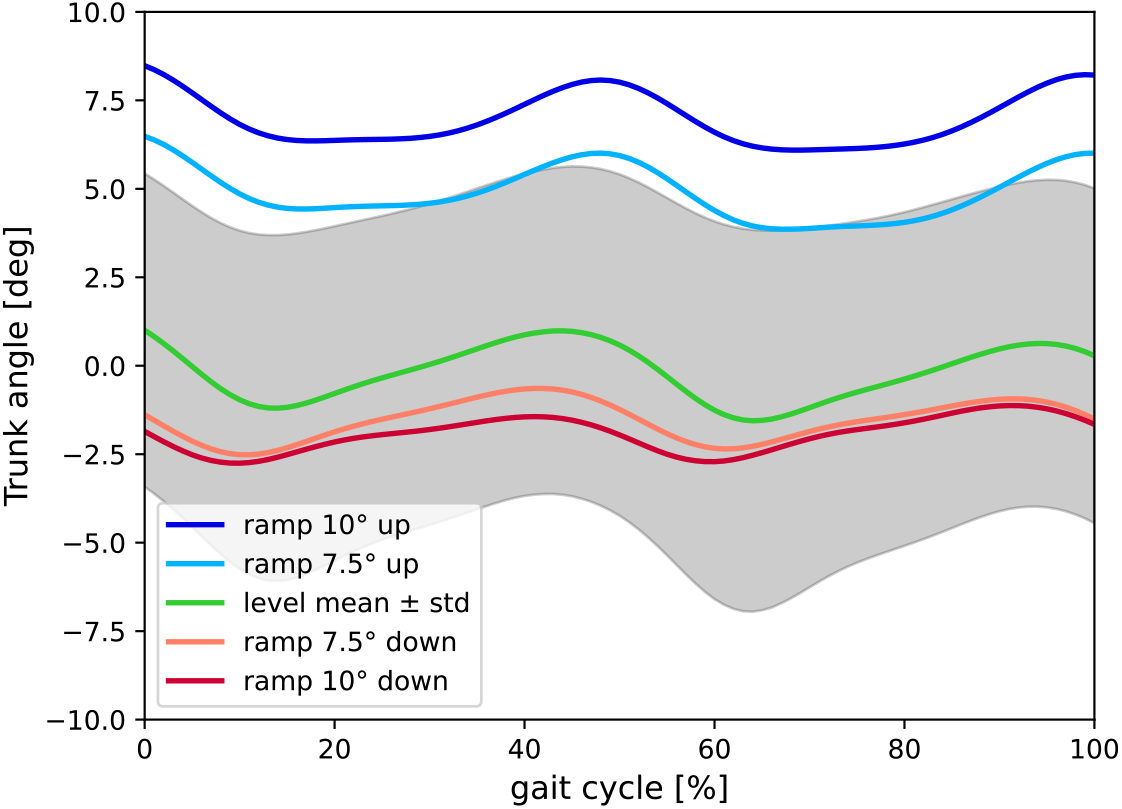
Trunk angle of human experiments. Mean values over all trials and participants are illustrated. For level walking, also standard deviation is shown (gray shaded area).

#### 3.1.1 VPP values

The mean value of the vertical VPP position (VPPz) ranges from 0.206 m (ramp 7.5° down) to 0.503 m (ramp 10° up). Significant differences can be found between level and upwards walking and between level and downwards walking of a ramp of 7.5° (Table 1). The horizontal VPP (VPPx) position ranges between −0.048 (ramp 7.5° up) and −0.032 (ramp 7.5° down) in mean values, that means they differ by <2cm. The single values are most around −0.04m ± 0.05 m (Figure 2). No significant differences between level and ramp walking could be observed. The *R*^2^-values are very high for all conditions (> 0.975), the mean value for level walking differs significantly from the mean values of upwards walking and from the mean value of walking down a ramp of 10°. In summary, walking up (down) shifts the VPP upwards (downwards), while *R*^2^ gets a bit lower (a bit lower, partially significant). This is also illustrated in the exemplarily VPP plots of one representative participant (Figure 2 b-d). Note that the VPP is calculated in a CoM-centered, trunk aligned coordinate frame.

#### 3.1.2 Joint moments

In Figure 3, joint moments and power for hip, knee and ankle joints are illustrated. During the first half of the gait cycle, the hip moment shows a slight increase during uphill walking and a slight decrease during downhill walking compared with level walking, whereas no differences are observed during the second half of the gait cycle. These changes are not statistically significant. Hip power increases during the stance phase in uphill walking and decreases in downhill walking relative to level walking. For the knee and ankle joints, more pronounced differences are observed, particularly during downhill walking. The knee extension moment increases significantly throughout the entire stance phase, accompanied by greater negative knee joint power. At the ankle, the moment increases during early stance and decreases during the push-off phase compared with level walking. During uphill walking, the knee extension moment increases in the first half of the stance phase and decreases in the second half, with the knee generating greater positive power than during level walking. The ankle moment remains comparable to level walking, while ankle power shows a significant increase during the push-off phase.

#### 3.1.3 Trunk angle

The behavior of the trunk angle in the experiments is illustrated in Figure 4. For level walking, the trunk angle is around zero (mean value is −0.2°), that means, the upper body is nearly upright. At walking down, backwards leaning can be observed (−1.9° for 10° ramp down), but no significant differences from level walking are measured. For walking up, the upper body is leaned more forwards significantly (6.9° for 10° ramp up). The trunk oscillation range for level walking is 2.6°. For walking up a ramp of 10°, the trunk oscillates by 2.4°, for walking down by 1.6°, respectively (Table 1).

### 3.2 Simulation Results

#### 3.2.1 VPP position

The simulations confirmed that the VPP systematically adapts to slope conditions (Fig. 2). On positive slopes, the VPP shifted anteriorly relative to the CoM, whereas on negative slopes it shifted posteriorly. Interestingly, the simulations revealed a linear coupling between vertical and horizontal VPP displacement. Specifically, an increase in backward/forward shift (VPPx) necessitated a proportional increase in VPP height (VPPz) in order to maintain a stable gait on a given slope.

#### 3.2.2 Hip torque

Hip torque modulation emerged as a key determinant of successful slope locomotion (Fig. 5). When the slope was set to level ground, increasing VPP height resulted in larger peak torques in both flexion and extension (see Fig. 5a). Under positive slope conditions, the model consistently predicted an increase in the magnitude of positive hip torque (see Fig. 5b). Conversely, for negative slopes, negative hip torque increased (see Fig. 5c). This change in the peak hip torque is more obvious when VPP height was fixed (see Fig. 5d). A systematic transition was observed across slope conditions: positive slopes amplified hip torque in the first half of the stance phase and reduced the hip torque in the second half, while negative slopes behaved in the opposite way (Fig. 5d).

**Figure 5.**
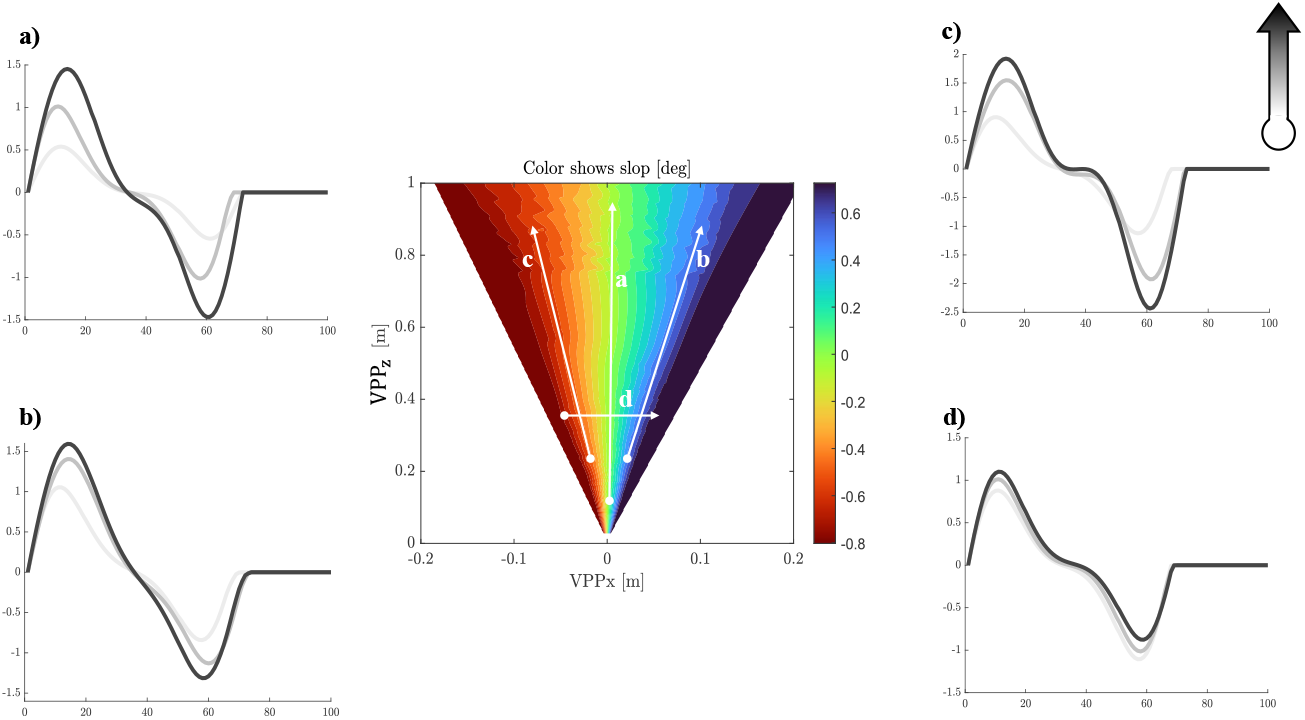
Simulation results: The central panel illustrates the virtual pivot point (VPP) position generated for different slope conditions. (a) Effect of increasing VPP height on hip torque during level walking. (b) Hip torque profiles on a positive slope for different VPP positions. (c) Hip torque profiles on a negative slope for different VPP positions. (d) Hip torque response when VPP height is fixed while both VPP horizontal position and slope vary systematically from negative to positive. Panels a–d are illustrated in the central subfigure, with movement direction color-coded (white to black indicating start to end of the arrow).

#### 3.2.3 Trunk orientation

The model also captured systematic adaptations in trunk kinematics (Fig. 6). On positive slopes, the trunk inclined forward, whereas on negative slopes it inclined backward. The degree of trunk lean scaled with slope angle, reflecting the need to realign the upper body to maintain balance and facilitate effective force transmission through the stance leg. VPP height had minimal influence on mean trunk inclination but strongly modulated trunk oscillation amplitude. Higher VPP positions were associated with larger oscillations.

**Figure 6.**
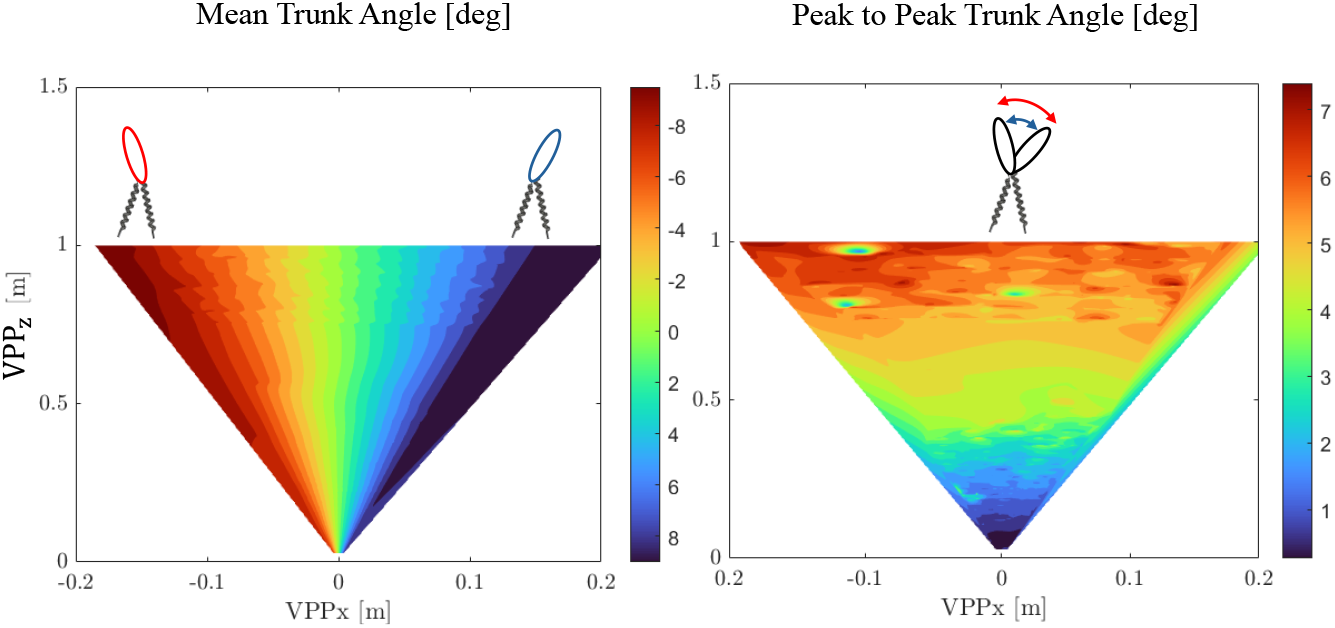
Simulation results: (a) Trunk inclination as a function of different virtual pivot point (VPP) positions. (b) Magnitude of trunk oscillations as a function of different VPP positions.

## 4 Discussion

Our integrated approach, combining template-model simulations with human experiments, reveals that the VPP is a critical component in the control of walking on a slope. The results demonstrate a systematic adaptation of VPP location and trunk dynamics in response to slope, providing a robust strategy for managing the energetic and stability demands of ramp walking. This discussion will focus on the role of the VPP, the complementary function of the leg axis, and the broader implications for understanding human locomotion.

### 4.1 VPP and Trunk Dynamics

First and foremost, our experimental results confirmed the existence of a Virtual Pivot Point during human walking on all tested slopes. The high coefficient of determination (*R*^2^ > 0.975) across all conditions indicates a strong convergence of the ground reaction forces, validating the VPP as a robust feature of gait, not only on level ground but also on inclines (see Table 1). This finding aligns with and extends a growing body of evidence establishing the VPP as a fundamental component of human locomotion across a variety of conditions, including level-ground walking [3, 24], running [8, 25], and across different walking styles, age groups, and even people with impaired gait [4, 5, 9, 26]. The consistent presence of the VPP highlights its importance as a core principle in the neural control of movement. Both the simulation and experimental results indicate that the VPP adapts systematically to the slope. Because the VPP represents the point where GRFs intersect in a coordinate frame attached to the CoM, its position directly influences whole-body angular momentum [4]. In the experiments, uphill walking produced a superior shift of the VPP accompanied by a forward trunk inclination, whereas downhill walking produced an inferior VPP shift with a slightly backward trunk lean. The simulation showed an anterior displacement of the VPP during uphill walking accompanied by a forward trunk inclination and a posterior displacement during downhill walking with a backward trunk lean. The amount of horizontal displacement also depends on the VPP height. Together, these observations align with prior findings that slope walking requires substantial adjustments in the regulation of whole-body angular momentum [14]. For instance, the forward trunk lean during uphill walking facilitates the generation of net positive work required to propel the center of mass upwards, while the backward lean during downhill walking is critical for controlled energy dissipation and braking to ensure stability [13, 14]. The FMCH controller in our model, which emulates the VPP concept, successfully reproduces these adaptations. By modulating the hip torque, the controller adjusts the ground reaction force to either reduce the braking component for uphill walking or increase it for downhill walking. This mechanism effectively manages the energy fluctuations introduced by the slope.

### 4.2 The Role of Leg Function in Large-Scale Energy Management

The divergence between our VPP-based model and empirical human walking data offers insight into a hierarchical control architecture in human locomotion. While our model remains stable on mild slopes, it fails to reproduce the robustness of human gait on steeper inclines and tends to adopt implausible trunk inclinations under high slope demands. This suggests that a pure VPP or hip-centric strategy, although a core element, is insufficient to account for the full adaptive repertoire of human walking [27, 28], especially beyond moderate slopes (e.g., > 8° in our experiments, compared to 1° in our simulation). In particular, our model required a pronounced horizontal shift of the VPP (anterior shift for uphill, posterior shift for downhill) compared to CoM, with the magnitude of the shift sensitive to the VPP height. In contrast, our experimental measurements showed no statistically significant horizontal VPP shift across slope conditions. This suggests that humans do not rely solely on hip torque guidance of the GRF toward the VPP, but instead engage a more distributed multi-joint control strategy.

In the human locomotor system, the ankle and knee joints provide further degrees of freedom to modulate the GRF direction and manage large mechanical energy fluctuations. In downhill (negative slope) walking, the leg joints increasingly assume an energy-absorption role. Our data show that the knee joint exhibits substantially higher extension torques during stance phase, functioning as a brake/damper to absorb excess energy. This is consistent with prior studies showing that knee extensors (e.g., quadriceps) contribute significantly to energy dissipation on declines [14, 29, 30]. Additionally, the ankle joint displays a significant torque increase during the first half of the stance phase, followed by a decrease in the second half in line with literature [29]. Although our simulation shows that using the hip mechanism by shifting VPP backward compared to CoM and bending the trunk backward, it is possible to navigate a negative slope, the human body’s ability to bend backward (trunk extension) is significantly limited due to several anatomical and biomechanical constraints [31] (see Fig. 4). Therefore, humans shift their strategy from the hip to the knee and ankle joint for navigating a negative slope. Looking at the joint’s power curve also shows that hip contribution compared to the knee and ankle is negligible for managing energy on a negative slope (see Fig. 3b).

In uphill walking, humans appear to adopt a coordinated, multi-joint strategy to realize the VPP. In the model, the leg is represented as a single spring, and control is implemented through a virtual hip torque defined between the upper body and a virtual leg—a line connecting the hip to the ankle. In humans, however, the leg is segmented; therefore, generating this virtual hip torque and effectively regulating the VPP requires coordinated control of both the hip and knee joints [32]. Correspondingly, our experimental data show significant changes in the hip and knee torque and power. Consistent with simulation predictions, hip torques play a major role in early stance (see Fig. 3a), and the trunk exhibits a forward inclination of about 7°, with the direction of tilt matching the model’s prediction. However, the model would predict a comparable trunk inclination only for slopes of approximately 1°, indicating that trunk dynamics and hip control alone cannot account for the observed adaptations [33]. Therefore, leg’s joints (knee, ankle) must contribute substantially to managing the energetic demands of steeper slopes.

In sum, the discrepancy between our VPP-centric model and human data underscores the necessity of a hierarchical, multi-joint control paradigm in human locomotion. The VPP or hip-strategy may afford stability and energy management on gentle slopes, but to robustly navigate steeper inclines or declines, humans recruit the knee and ankle to share the load, provide dissipation or propulsion as needed, and ensure stable gait without adopting mechanically implausible postures.

### 4.3 Trunk Oscillation, VPP Height, and Energy Management

In our simulations, increasing the VPP height in the sagittal plane led to greater trunk oscillations, which parallels our experimental observation that uphill walking is characterized by a higher VPP and larger trunk oscillations, while downhill walking shows a lower VPP and smaller oscillations. When shifting VPP upward, we are increasing the length of the virtual pendulum and also increasing the lever arm of the GRF around CoM. This means that the oscillation of the whole-body angular momentum is increasing. These findings are consistent with Silverman et al., who reported that the range of whole-body angular momentum is larger on inclines but similar or reduced on declines compared to level walking, particularly in the sagittal and frontal planes [14].

Functionally, this asymmetry reflects a slope-dependent trade-off between propulsion and stability. The regulation of angular momentum has previously been shown to be important in fall prevention [34]. It appears that actively controlling angular momentum to a greater extent (i.e., reducing the range of angular momentum) while walking down a decline is a protective mechanism to prevent slipping or falling. Such tight control may not be necessary for walking up an incline, when the risk of falling is not as high [35]. Modulating the VPP pendulum length—which increases with VPP height and defines the moment of inertia of the pendulum—allows the system to potentially store more energy in this degree of freedom. Therefore, the modulation of VPP pendulum length could serve as a tool to insert or withdraw energy from the system, influencing energy management. It is also shown that a moderate oscillation in angular momentum is energy effitient [36].

Together, these findings support a control strategy in which the trunk acts as an actively tuned oscillator: its motion is amplified during uphill walking to assist energy redistribution, but constrained during downhill walking to maintain stability while the lower limbs assume the primary role of energy dissipation. This slope-dependent modulation of trunk oscillations, in conjunction with changes in VPP height, provides a unified framework for understanding how humans balance the competing demands of propulsion and stability in sloped locomotion.

### 4.4 Implications for Assistive Devices

The principles of locomotor control elucidated in this study have direct implications for the next generation of assistive devices, including wearable exosuits, exoskeletons, and robotic prostheses. The FMCH-based hip torque modulation strategy, which emulates the VPP principle, has already been employed to control assistive devices [24, 32, 37]. Our results extend this framework by demonstrating that VPP dynamics adapt systematically to non-level terrains: the VPP shifts upward during uphill walking and downward during downhill walking. This suggests that incorporating slope-adaptive VPP modulation could achieve both energy efficiency and balance (see Sec. 4.3). The FMCH controller is particularly attractive for device control because of its low-dimensional parameterization. It can be described by two main parameters: a gain term, which primarily regulates the VPP height, and a rest angle, which adjusts the horizontal alignment of the VPP relative to the CoM. Adjusting these two parameters allows the controller of the assistive device to potentially tune the VPP position across different terrains, thereby modulating whole-body angular momentum and trunk dynamics in a bioinspired manner [38].

### 4.5 Limitations and Future Research

Several limitations of this study should be considered when interpreting the results. First, our analysis was restricted to the sagittal plane. While this approach captures the primary direction of locomotor dynamics during slope walking, it neglects adaptations in the frontal plane. Previous studies have demonstrated that frontal-plane mechanics and angular momentum regulation also change systematically with slope, contributing to mediolateral balance and stability control [29, 30, 39]. A full three-dimensional analysis would therefore provide a more complete picture of how humans modulate VPP and trunk dynamics across complex terrains. Second, our template model employed a leg with fixed stiffness. This simplification limited the capacity to interpret the role of the leg in energy redistribution and joint coordination. Although a fixed-stiffness leg was sufficient for our focus on VPP dynamics, it constrains the generalizability of our findings regarding detailed leg function. Finally, our participant sample consisted of healthy young adults, and slopes were limited to ± 7.5° and ± 10° on a controlled ramp environment. This constrains the generalizability of our findings to flatter or steeper terrains, or populations with altered locomotor control (e.g., older adults or clinical cohorts) [4]. Extending future work to more diverse populations will be critical to fully assess the role of VPP dynamics in human walking.

## Notes

### Competing Interest Statement

The authors have declared no competing interest.

### Funding

- Deutsche Forschungsgemeinschaft: 450821862

## References

1. Arthur H Dewolf, Yuri P Ivanenko, Francesco Lacquaniti, and Patrick A Willems. Pendular energy transduction within the step during human walking on slopes at different speeds. PLoS One, 12(10):e0186963, 2017.

2. Hermann Schwameder. Moving on slopes: issues and challenges from a biomechanical perspective. ISBS Proceedings Archive, 36(1):4, 2018.

3. H-M Maus, SW Lipfert, M Gross, J Rummel, and A Seyfarth. Upright human gait did not provide a major mechanical challenge for our ancestors. Nature communications, 1(1):70, 2010.

4. Vahid Firouzi, Omid Mohseni, Andre Seyfarth, Oskar von Stryk, and Maziar A Sharbafi. Exploring the control of whole-body angular momentum in young and elderly based on the virtual pivot point concept. Royal Society Open Science, 11(9):240273, 2024.

5. Johanna Vielemeyer, Lucas Schreff, Stefan Hochstein, and Roy Müller. Virtual pivot point: Always experimentally observed in human walking? Plos one, 18(10):e0292874, 2023.

6. Johanna Vielemeyer, Nora-Sophie Staufenberg, Lucas Schreff, Daniel Rixen, and Roy Müller. Walking like a robot: do the ground reaction forces still intersect near one point when humans imitate a humanoid robot? Royal Society Open Science, 10(5):221473, 2023.

7. Roy Müller, Christian Rode, Soran Aminiaghdam, Johanna Vielemeyer, and Reinhard Blickhan. Force direction patterns promote whole body stability even in hip-flexed walking, but not upper body stability in human upright walking. Proceedings of the Royal Society A: Mathematical, Physical and Engineering Sciences, 473(2207):20170404, 2017.

8. Vahid Firouzi, Fariba Bahrami, and Maziar A Sharbafi. Human balance control in 3d running based on virtual pivot point concept. Journal of Experimental Biology, 225(4):jeb243080, 2022.

9. Patrick Scholl, Vahid Firouzi, Mohammad Taghi Karimi, Andre Seyfarth, and Maziar A Sharbafi. Virtual pivot point model predicts instability in parkinsonian gaits. In 2023 IEEE International Conference on Systems, Man, and Cybernetics (SMC), pages 5261–5266. IEEE, 2023.

10. Nathaniel T Pickle, Alena M Grabowski, Arick G Auyang, and Anne K Silverman. The functional roles of muscles during sloped walking. Journal of biomechanics, 49(14):3244–3251, 2016.

11. Andrea N Lay, Chris J Hass, D Webb Smith, and Robert J Gregor. Characterization of a system for studying human gait during slope walking. Journal of applied biomechanics, 21(2):153–166, 2005.

12. Andrew H Hansen, Dudley S Childress, and Steve C Miff. Roll-over characteristics of human walking on inclined surfaces. Human movement science, 23(6):807–821, 2004.

13. Amy Silder, Thor Besier, and Scott L Delp. Predicting the metabolic cost of incline walking from muscle activity and walking mechanics. Journal of biomechanics, 45(10):1842–1849, 2012.

14. Anne K Silverman, Jason M Wilken, Emily H Sinitski, and Richard R Neptune. Whole-body angular momentum in incline and decline walking. Journal of biomechanics, 45(6):965–971, 2012.

15. Plug-in Gait Reference Guide. https://help.vicon.com/space/Nexus216/11607059/Plug-in+Gait+Reference+Guide. Accessed: 2025-10-07.

16. Herman J Woltring. A fortran package for generalized, cross-validatory spline smoothing and differentiation. Advances in Engineering Software (1978), 8(2):104–113, 1986.

17. Hugh Herr and Marko Popovic. Angular momentum in human walking. Journal of experimental biology, 211(4):467–481, 2008.

18. John D Storey. False discovery rate. In International encyclopedia of statistical science, pages 944–949. Springer, 2025.

19. Johanna Vielemeyer, Lisa Tronicke, Lucas Schreff, Rainer Abel, Knut Lechler, and Roy Müller. A full-body motion capture gait dataset of healthy young adults walking ramps up and down. Scientific Data, 2026.

20. Mohammad Shahbazi, Robert Babuška, and Gabriel AD Lopes. Unified modeling and control of walking and running on the spring-loaded inverted pendulum. IEEE Transactions on Robotics, 32(5):1178–1195, 2016.

21. Maziar A Sharbafi and Andre Seyfarth. Fmch: A new model for human-like postural control in walking. In 2015 IEEE/RSJ International Conference on Intelligent Robots and Systems (IROS), pages 5742–5747. IEEE, 2015.

22. Johanna Vielemeyer, Eric Grießbach, and Roy Müller. Ground reaction forces intersect above the center of mass even when walking down visible and camouflaged curbs. Journal of experimental biology, 222(14):jeb204305, 2019.

23. Jasper Snoek, Hugo Larochelle, and Ryan P Adams. Practical bayesian optimization of machine learning algorithms. Advances in neural information processing systems, 25, 2012.

24. Vahid Firouzi, Omid Mohseni, and Maziar A Sharbafi. Model-based control for gait assistance in the frontal plane. In 2022 9th IEEE RAS/EMBS International Conference for Biomedical Robotics and Biomechatronics (BioRob), pages 1–8. IEEE, 2022.

25. Özge Drama, Johanna Vielemeyer, Alexander Badri-Spröwitz, and Roy Müller. Postural stability in human running with step-down perturbations: an experimental and numerical study. Royal Society open science, 7(11):200570, 2020.

26. Johanna Vielemeyer, Cristina Sole, Manuela Galli, Matteo Zago, Roy Müller, and Claudia Condoluci. A study on the intersection of ground reaction forces during overground walking in down syndrome: effects of the pathology and left–right asymmetry. Symmetry, 15(2):544, 2023.

27. Jana R Montgomery and Alena M Grabowski. The contributions of ankle, knee and hip joint work to individual leg work change during uphill and downhill walking over a range of speeds. Royal Society open science, 5(8):180550, 2018.

28. M Kuster, S Sakurai, and GA Wood. Kinematic and kinetic comparison of downhill and level walking. Clinical biomechanics, 10(2):79–84, 1995.

29. Sang-Kyoon Park, Hyun-Min Jeon, Wing-Kai Lam, Darren Stefanyshyn, and Jiseon Ryu. The effects of downhill slope on kinematics and kinetics of the lower extremity joints during running. Gait & posture, 68:181–186, 2019.

30. Nathalie Alexander and Hermann Schwameder. Lower limb joint forces during walking on the level and slopes at different inclinations. Gait & posture, 45:137–142, 2016.

31. Michael A Adams, Stephen May, Brian JC Freeman, Helen P Morrison, and Patricia Dolan. Effects of backward bending on lumbar intervertebral discs: relevance to physical therapy treatments for low back pain. Spine, 25(4):431–438, 2000.

32. Vahid Firouzi, Ayoob Davoodi, Fariba Bahrami, and Maziar A Sharbafi. From a biological template model to gait assistance with an exosuit. Bioinspiration & biomimetics, 16(6):066024, 2021.

33. Richard W Nuckols, Kota Z Takahashi, Dominic J Farris, Sarai Mizrachi, Raziel Riemer, and Gregory S Sawicki. Mechanics of walking and running up and downhill: A joint-level perspective to guide design of lower-limb exoskeletons. PloS one, 15(8):e0231996, 2020.

34. Mirjam Pijnappels, Maarten F Bobbert, and Jaap H Van Dieën. Push-off reactions in recovery after tripping discriminate young subjects, older non-fallers and older fallers. Gait & posture, 21(4):388–394, 2005.

35. Mark S Redfern, Rakié Cham Krystyna Gielo-Perczak, Raoul Grönqvist, Mikko Hirvonen, Håkan Lanshammar, Mark Marpet, Clive Yi-Chung Pai IV, and Christopher Powers. Biomechanics of slips. Ergonomics, 44(13):1138–1166, 2001.

36. Myriam L De Graaf, Juul Hubert, Han Houdijk, and Sjoerd M Bruijn. Influence of arm swing on cost of transport during walking. Biology open, 8(6):bio039263, 2019.

37. Arjang Ahmadi, Vahid Firouzi, Dennis Haufe, Sebastian Hirt, Andre Seyfarth, Gregory S Sawicki, Rolf Findeisen, and Maziar Ahmad Sharbafi. Towards real-time personalized control in wearable robotics: A hierarchical architecture for lower-limb assistance. In 2025 IEEE 64th Conference on Decision and Control (CDC), pages 5411–5418. IEEE, 2025.

38. Omid Mohseni, Serajeddin Ebrahimian, Vahid Firouzi, Morteza Khosrotabar, Mario Kupnik, Maziar A Sharbafi, and André Seyfarth. Exploring the virtual pivot point in unilateral transfemoral amputee locomotion: Implications for prosthetic development. In 2025 IEEE/RSJ International Conference on Intelligent Robots and Systems (IROS), pages 8254–8260. IEEE, 2025.

39. K Masayuki, Kazutaka Hata, Ryoji Kiyama, Tetsuo Maeda, and Kazunori Yone. Biomechanical characterization of slope walking using musculoskeletal model simulation. Acta of bioengineering and biomechanics, 20(1), 2018.

